# The superior colliculus gates dopamine responses to conditioned stimuli in visual classical conditioning

**DOI:** 10.1101/2025.01.06.631486

**Authors:** Yan-Feng Zhang, Jean-Philippe Dufour, Peter Zatka-Haas, Peter Redgrave, Melony J Black, Armin Lak, Ed Mann, Stephanie J Cragg, Wickliffe C Abraham, John NJ Reynolds

**Author notes:** Correspondence to: Dr Yan-Feng Zhang, Prof John N. J. Reynolds.

## Abstract

In classical Pavlovian conditioning, it is well-established that midbrain dopamine neurons respond to conditioned stimuli (CS) that predict a reward. However, how the dopamine neurons associate a neutral CS to a reward remains unknown. Here, we show that the superior colliculus (SC) develops neuronal responses to a visual CS during conditioning, which in turn drive the responses of dopamine neurons. Visual responses in the SC were only potentiated when a behaviorally meaningful time interval separated the visual stimulus and reward. Potentiation also required the convergence of visual, dopamine and serotonin inputs to the SC. Importantly, blocking potentiation of the visual response was sufficient to suppress the dopamine responses following a CS. These results reveal a mechanism for how the brain forms associations between unconditioned stimuli and behaviorally meaningful visual information during classical conditioning.

## Introduction

The ability to identify salient events as predictive of subsequent reward or danger is critical for survival. Classical conditioning, described by Pavlov more than a century ago^1^, is believed to be underpinned by brain mechanisms that associate a neutral sensory event with a reward. Much evidence implicates the phasic activity of midbrain dopamine neurons following a sensory event as reflecting processes of classical conditioning^2, 3^. Thus, after repeated pairing with an unconditioned appetitive stimulus (US), a neutral stimulus becomes conditioned, and the dopamine neurons in midbrain fire burst activity following the conditioned stimulus (CS) to reflect that a reward is predicted^2, 4^. For example, in visual classical conditioning, dopamine neurons start to fire in bursts at about 100 ms after a visual cue is applied to the eyes, and this dopamine response is critical for animals to learn the salience of the visual cues. But where does the formation of the US-CS association take place? The midbrain dopamine neurons do not receive primary visual input. In addition, despite the marked heterogeneity in dopamine neuron firing patterns^5, 6^, dopamine neurons show a unified response to visual stimuli at individual cell bodies^7, 8^ in visual classical conditioning. Thus, it is possible that dopamine neurons receive input from another brain area that itself develops the conditioned response to a visual CS.

To be the upstream brain structure that drives the dopamine response to a CS, a candidate brain structure should display a conditioned response that correlates with the presence or absence of reward. Specifically, this response should increase when the CS is associated with the US and reduce when the US is removed. Additionally, these conditioned responses should occur consistently at a shorter latency than the dopamine neuron responses, with a latency difference suitable for the candidate brain structure to convey its output to the dopamine neurons. Furthermore, the conditions required to generate this activity change should match those that generate the dopamine response. Finally, if the response to the CS contributes to the dopamine phasic activity, blocking the cellular plasticity in the candidate structure should be sufficient to decrease dopamine neuron responsiveness to the CS.

Here, we hypothesise that the superior colliculus (SC), a primary sensory area in the midbrain, is the brain area that identifies the salience of visual cues in classical conditioning and drives dopamine phasic activity. The superficial layers of the SC receive visual input from the retina and relay it to the deep layers, which have direct glutamatergic projections to midbrain dopamine neurons^9^. When dis-habituated pharmacologically, the visual input through this pathway can drive midbrain dopamine neurons at a latency similar to that observed in classical conditioning^10^. By using *in vivo* field potential recording, multi-unit recording, fast-scan cyclic voltammetry, and calcium imaging in anaesthetised and freely moving animals, we show that the deep layers of SC respond to a visual CS similarly to dopamine neurons but faster with a consistent short latency. We propose this response to be the main trigger of dopamine neuron burst activity following a visual CS in classical conditioning.

## Results

### SC responds to conditioned visual stimuli like dopamine neurons but consistently faster

For the SC to be the region where visual stimuli are associated with reward, it should develop a response to reward-associated visual stimuli that decreases during extinction. To test this, a flash of light into the eye was used as the CS, electrical stimulation of the SNc/VTA was used as the rewarding US ^11, 12^ and the associated visual evoked potential (VEP) was measured in the deep layers of the contralateral SC in anaesthetised rats (**Fig. 1a, Extended Data Fig. 1**). Stimulation of SNc/VTA was applied 1 s following the light flash to mimic a physiologically meaningful reward delay. The paired visual and SNc/VTA stimulation was applied for 60 pairings. Compared to the baseline, a negative deflection of the VEP (nVEP) developed (84.5 ± 7.8 µV; mean ± SEM) with a peak latency of ∼70 ms (**Fig. 1b,c**).

**Fig. 1.**
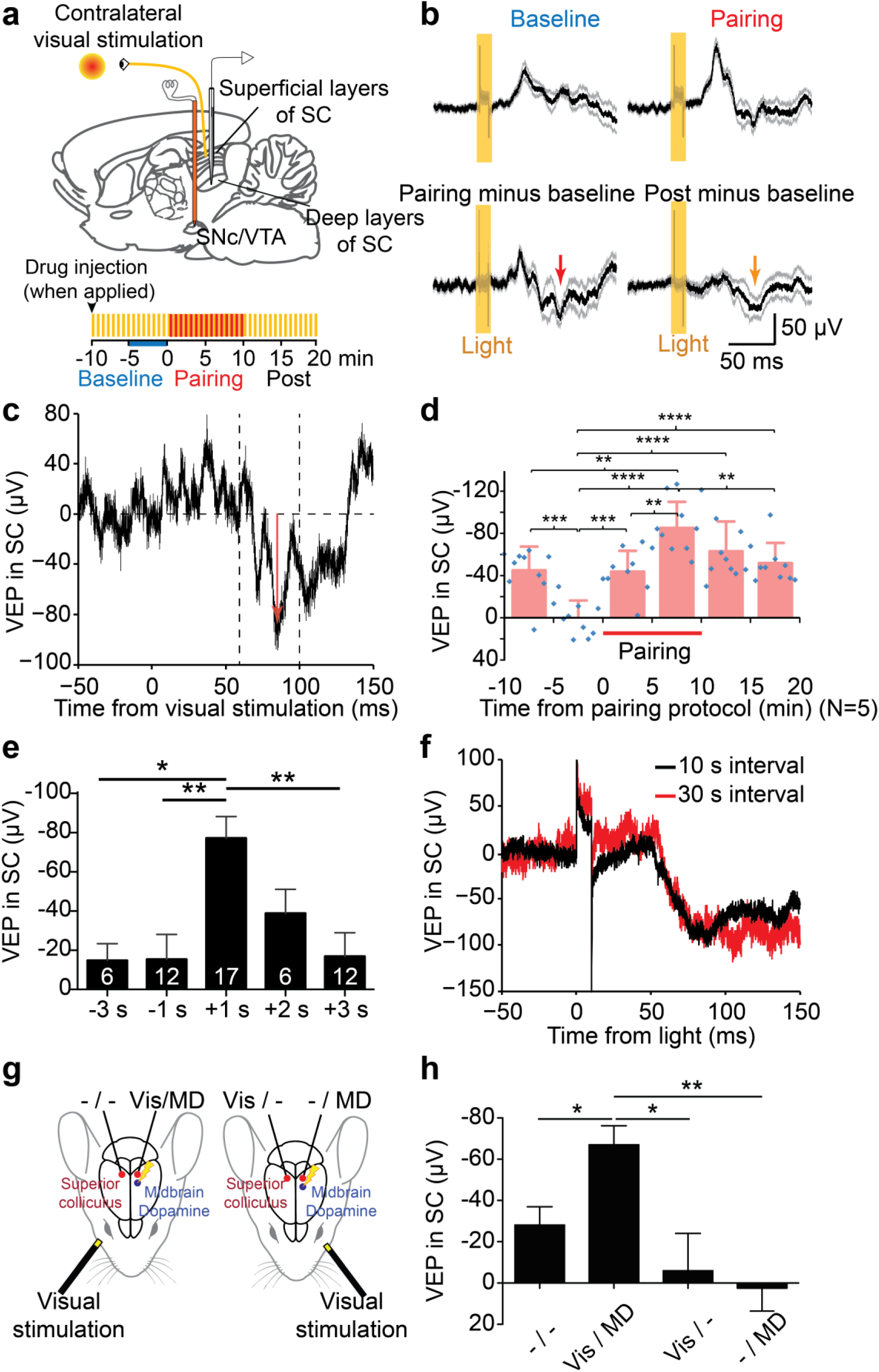
Potentiation of the negative VEP component (nVEP) in the deep layers of the SC. **a**, Cartoon of the experimental setup on a sagittal rat brain section. Lower panel, a diagram of the experimental protocol with timing of light flashes alone (orange) and light flashes paired with SNc/VTA stimulation (red). **b**, Effect of a light flash on SC responses (S.D., grey traces) before and after SNc/VTA pairing (N = 9 animals). **c**, Representative VEP from one animal showing the measured negative component (nVEP; arrowed). **d**, The mean nVEP (N = 5 animals) plotted across time throughout the whole experiment (blue dots; histograms 5-min bins, inverted so negative is up; One-way ANOVA with Tukey’s Multiple Comparison Test). **e**, The nVEP is significantly potentiated only when light is applied 1 s before the SNc/VTA stimulation reward (One-way ANOVA with Bonferroni’s Multiple Comparison Test). **f**, The nVEP was induced regardless of whether an inter-pairing interval of 10 s or 30 s was applied in the same experiment. Recording at the 30 s interval was commenced after the nVEP had returned to baseline levels following an inter-pairing interval of 10 s. **g**, Experimental layout from top of the animal’s head: the VEPs in both SCs (red circles) were recorded simultaneously. Each eye (black semi-circles) was stimulated alternately, and midbrain dopamine neurons (MD) on the left (blue circles) were stimulated following each light flash. **h**, A significant nVEP component only developed when visual input and midbrain dopamine stimulation spatially converged (Vis/ MD group; N = 8 animals). Friedman test (repeated, nonparametric) with Dunns post hoc test. Data are means ± S.E.M. * P < 0.05, ** P < 0.01, *** P < 0.001, **** P < 0.0001.

Closer inspection of the VEPs throughout the experimental protocol (**Fig. 1d**; N = 5 animals) revealed that nVEP not only gradually potentiated when visual stimulation was paired with reward but also then decreased during extinction, i.e. when the visual stimulus was disassociated from the reward, over the 10 min following pairing. Also, consistent with previous reports^14^, response to the initial visual stimulation habituated quickly during baseline testing (-10 to 0 min). There was a trend for nVEP extinction to be slower compared with the initial habituation to baseline light flashes (**Fig. 1d**; *P* = 0.056, t = 2.67, df = 4). This might reflect a higher salience value encoded in the visual stimulus after it is paired with a reward compared with when it is novel^15, 16^. Therefore, the amplitude and persistence of the nVEP in the deep SC was positively correlated with dopamine neuron activity during novel visual stimulation, as described in reinforcement learning and the ‘shaping bonuses’ theory, respectively ^17^.

### Critical temporal and spatial afferent convergence

We then tested whether the latency of the nVEP induced by classical conditioning was appropriate to trigger the dopamine response following a CS. It has been shown that local injection of bicuculline (BIC), a GABA_A_ receptor antagonist, into the SC in an anaesthetised animal disinhibit the SC, leading the deep layers of SC to respond to the visual stimulation with a nVEP and a response in midbrain dopamine neurons at a latency that is similar to classical conditioning ^10, 18^. Here we found that the nVEP induced by SNc/VTA pairing showed an onset latency similar to the nVEP induced by BIC (≈ 60 ms), although with smaller amplitude (80 µV pairing; 330 µV post-BIC) and earlier peak latency (80 ms, and 100 ms respectively; **Extended Data Fig. 2**). Therefore, the timing of the nVEP generated in SC is ideal for driving the dopamine response during classical conditioning. This also suggests that the mechanism underlying the pairing-induced potentiation of the nVEP may involve, in part, a reduction in inhibition within the collicular microcircuit.

Classical conditioning is optimal at particular inter-stimulus-intervals between CS and US ^19^. To test if this is true for the potentiated nVEP, we applied SNc/VTA stimulation at various times before or after visual stimulation (**Fig. 1e**). The nVEP was significantly larger when SNc/VTA stimulation was applied 1 s after the light compared to 3 s before, 1 s before, or 3 s after. Pairing at 2 s induced an intermediate potentiation. The potentiated response following 1 s pairing was insensitive to pairing frequency, with a similar potentiated response resulting for inter-pairing intervals of 10 s or 30 s (**Fig. 1f**). Therefore, nVEP potentiation can be induced through low pairing frequencies, with each pairing occurring within a behaviorally-relevant critical time window, consistent with behavioural observations in classical conditioning ^19^.

The brain structure that develops the response to the CS should receive information from both the CS and US to identify their association. To test whether the potentiated nVEP in SC fits this requirement, we alternated the visual stimulus between the two eyes while activating the midbrain dopamine neurons on the left side only (**Fig. 1g**). In rats, more than 90% of visual input to the SC is contralateral ^20^, and almost all projections from the SNc and VTA are restricted to the same hemisphere ^21^. We found that only the nVEPs in the SC where there was convergence of visual input (from contralateral side) and midbrain dopamine neuron activation (from the ipsilateral side) during pairing were potentiated (**Fig. 1h**). None of the other combinations of light flash and midbrain dopamine neuron stimulation elicited a comparable visual response. Thus, potentiation of the nVEP required a spatial convergence within the same hemisphere of activation of the primary sensory area in the SC and midbrain dopamine neuron activation.

### The nVEPs represent local spike activity in deep layers of the SC

We further confirmed that the nVEP in the deep layers of the SC represents local neuronal spike activity. The negative deflection of the local field potential has long been used as a proxy for excitatory input and local spike activity in brain areas such as the hippocampus^13^. To determine if nVEP is correlated to local neuronal activity in the SC, we recorded spike activity and nVEP simultaneously in the mouse SC using silicon probes (**Fig. 2a**).

**Fig. 2.**
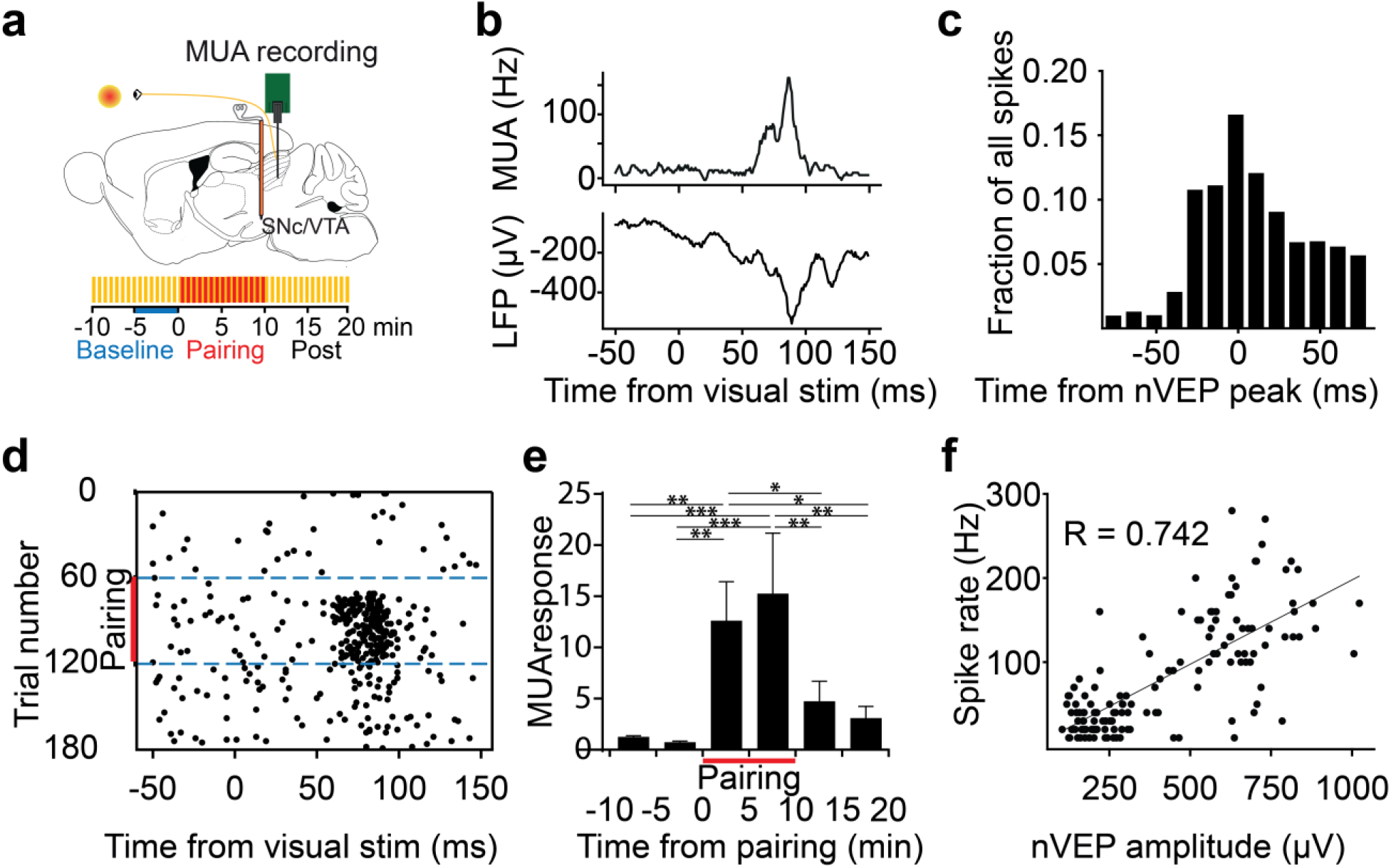
Amplitudes of nVEPs represent the number of action potentials generated from neurons in the deep layers of the SC. **a**, Cartoon of the experimental setup. **b**, Example average multi-unit activity (MUA) trace during pairing and LFP trace (pairing minus baseline) in a representative mouse, showing the nVEP peak coinciding with the MUA peak. **c**, Histogram of spikes during pairing across all responsive channels (n = 2926 spikes in 7 channels) and their time from the nVEP peak in each trial. **d**, Example raster plot of the MUA response to light stimulus in a representative channel. **e**, MUA response magnitude time course in all responsive channels as a fraction of the mean pre-pairing baseline response in each channel. (One-way ANOVA with Fisher’s LSD post hoc test, data are Mean ± S.E.M. * p < 0.05, ** p<0.001, *** p<0.0001) **f**, Scatter plot of nVEP amplitude versus MUA response spike rate for each trial in an example channel across a complete experiment.

First, we successfully replicated our findings of nVEP in the mouse SC, where nVEPs with similar latency and comparable amplitude were observed when neutral visual stimuli were paired with electrical stimulation of the mouse SNc/VTA. Additionally, the local action potentials and nVEP matched well in latency and magnitude (**Fig. 2b & c**).

We then found that the firing of local action potentials developed similarly to the nVEP during pairing, increasing during pairing and reducing when electrical stimulation was removed (**Fig. 2 d & e**). Furthermore, the number of action potentials recorded in the deep layers of the SC was highly correlated with the amplitude of the nVEP recorded simultaneously **(Fig. 2f)**.

These results suggest that the nVEP recorded in the deep layers of the SC is indeed associated with local spike activity.

### Rewarding midbrain dopamine stimulation also induces nVEP potentiation

To determine whether potentiation of the SC response could be induced using known behaviorally rewarding stimulation, stimulating electrodes were implanted in the SNc/VTA in eight rats, and the animals were operantly conditioned using an intracranial self-stimulation (ICSS) protocol^12^. Animals were then anesthetised, and the VEP was recorded using the same stimulus protocol, and the stimulus current was identified for each animal to be behaviorally rewarding^12^ (**Fig. 3a**,**b**). The rewarding current potentiated the nVEP in the SC in ICSS rats by a magnitude similar to that observed in naïve rats (**Fig. 3b**).

**Fig. 3.**
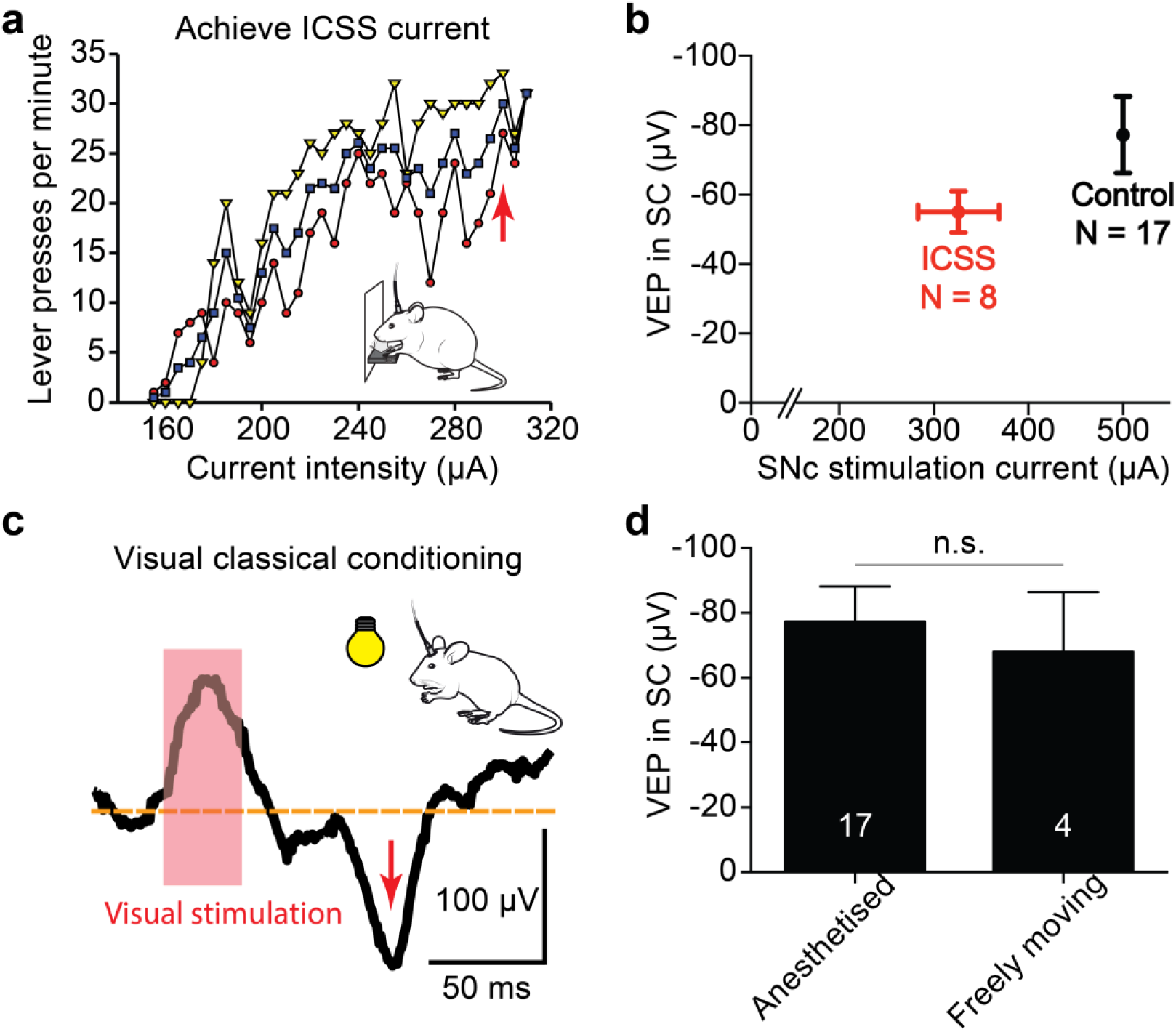
The nVEP component could be induced in both anaesthetised and freely moving rats, the latter receiving rewarding ICSS currents. **a**, Lever-pressing rate for one rat in response to increments (red circles) and decrements (yellow diamonds) in SNc/VTA stimulus intensity. Arrow indicates the optimal current that maximises the average rate (blue squares). **b**, nVEPs in anesthetised trained rats induced using optimal ICSS currents and 10 ms visual stimuli were comparable to those obtained in naïve rats using a standard current of 500 µA (unpaired t-test). **c**, An example VEP trace recorded from a freely moving rat (rat 1004), showing the nVEP component (arrow) following 1 second pairing between visual stimulation (30 ms; red bar) and ICSS-like stimulation. **d**, nVEPs were of similar amplitude in both anesthetised naïve rats and freely moving rats (unpaired t-test).

We then tested whether the nVEP in the SC would be potentiated during classical conditioning in awake animals. We first determined the intensity of the rewarding electrical stimulation on SNc/VTA by using an operant ICSS protocol^12^. Then, freely moving rats were placed in a chamber for visual classical conditioning. A bright whole-field visual stimulus was paired with their ICSS reward currents and the nVEP was recorded in the SC in response to the visual stimulus (**Fig. 3c**). We found the same potentiated SC response in these freely moving animals (68.0 ± 18.5 µV, N = 4) as recorded in the anesthetised rats (**Fig. 3d**). Moreover, following pairing, animals responded to the whole-field light flash with movement, consistent with others’ observations in classical conditioning^22^. This contrasted with the absence of such movements in response to the light during baseline or prolonged extinction (> 10 min), suggesting they had learnt through pairing the significance of the light flash as a predictor of SNc/VTA reward delivery. The change in the animal’s activity, calculated as the difference in speed of movements 0.5 s after compared to 0.5 s before the light flash, correlated with the amplitude of the potentiated VEP (r = 0.98, *P* < 0.05). In contrast, the change in movements in response to the midbrain dopamine stimulation did not correlate with the nVEP (r = 0.09, *P* = 0.91), indicating that potentiation of the nVEP was related to the conditioned change in response speed rather than the reward delivery (**Extended Data Fig. 3**). Therefore, potentiation of visual responses in the deep layers of the SC was induced in behaving animals and was associated with acquired responsiveness to a visual CS.

### Neuromodulator role in the potentiation of the SC neuronal response

In order to investigate the cellular mechanisms underlying the potentiation of SC VEPs, antagonists to dopamine or serotonin (5HT) receptors were locally injected into the SC immediately before the commencement of the light flash baseline (**Fig. 4a**). Both the D_1_-like dopamine receptor antagonist SCH23390 (17.8 ± 17.3 µV) and the 5-HT_1A_ receptor antagonist WAY100635 (-36.8 ± 19.4 µV) significantly blocked the induction of the potentiated nVEP component compared to the control (uninjected) group (**Fig. 4b&c**). Local saline injection (69.1 ± 8.8 µV) did not affect the potentiation of the nVEP compared to the control (uninjected) group (77.2 ± 11.1 µV), but was significantly different to the SCH23390 and WAY100635 groups. In contrast, the 5-HT_2A_ receptor antagonist ketanserin did not affect the induction of the nVEP component (66.3 ± 12.3 µV) when compared to either normal or saline controls (**Fig. 4b**).

**Fig. 4.**
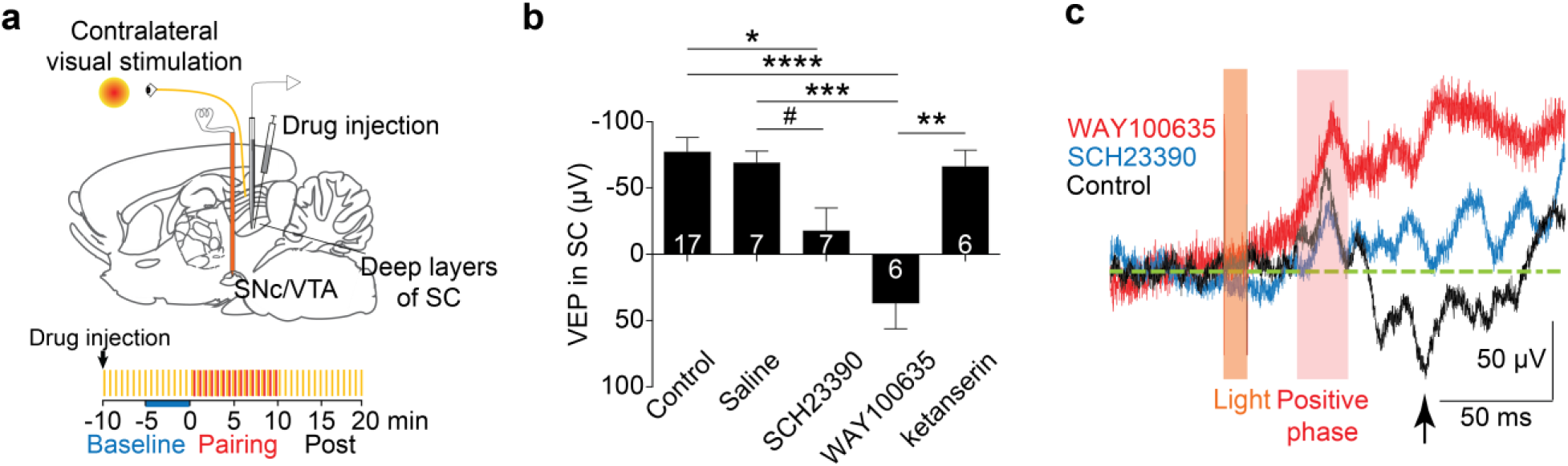
Potentiation of the nVEP component required dopamine D_1_ and serotonin 5-HT_1A_ but not 5-HT_2A_ receptor activation within SC. **a**, Cartoon of the experimental setup on a sagittal rat brain section with indication of local drug injection into the deep layers of SC. **b**,The nVEP in the deep layers of the SC was blocked by local injection of SCH23390 (D1 antagonist) and WAY100635 (5-HT1A antagonist) but not saline or ketanserin (5-HT2A antagonist). One-way ANOVA with Bonferroni’s Multiple Comparison Test. # p<0.05 Paired t-test. **c**, The control, SCH23390 and WAY100635 groups all showed similar positive VEP components (light red shaded area), while only control experiments showed an enhanced negative component (arrow).

Collectively, these results suggest that potentiation of the nVEP in the deep layers of the SC is mediated by local dopamine acting on D_1_ receptors and serotonin acting on 5-HT_1A_ but not 5- HT_2A_ receptors. This is the first functional evidence of dopamine and serotonin in the deep layers of the SC and their role in potentiation of visual responses.

### US and CS is associated in the deep layers of SC

We then investigated whether the deep layers of the SC are the site of generation of the response to the CS, or whether the response is driven by the upstream superficial layers of the SC. Of note, the positive deflection of the VEP (pVEP) in the deep layers of the SC, which reflects the neuronal activity in the superficial layers (**Extended Data Fig. 4**), was also potentiated during the pairing protocol (≈ 30 to 40 ms latency; see **Fig. 1c**), but this effect did not persist during the extinction period, unlike the negative component (**Fig. 1b**; pairing minus baseline c.f. post minus baseline). This indicates that the potentiated nVEP in the deep layers is not secondary to the potentiation of neuronal activity in the superficial layers. In addition, SCH23390 or WAY100635 diminished only the nVEP but not the pVEP during CS-US pairing (**Fig. 4b**), indicating that different cellular mechanisms are operating in superficial and deep layers of the SC. Taken together, our data strongly suggest that the potentiation of the nVEP representing the conditioned visual stimulus originated within the intermediate/deep layers of the SC.

### Potentiation of visual responses in SC drives striatal dopamine release

Finally, we tested whether conditioning of the nVEP in the SC is sufficient to drive midbrain dopamine neurons to release dopamine in target areas during visual classical conditioning. Using in vivo FCV, we monitored extracellular striatal dopamine in the ventral striatum of anesthetised adult C57Bl6/J mice (**Fig. 5a**). To amplify dopamine signals, cocaine (20 mg/kg), a dopamine reuptake inhibitor, was injected intraperitoneally. The same pairing protocol that potentiated the nVEP in the SC induced a new striatal dopamine signal in response to the light, which had been absent at baseline (**Fig. 5b**). To confirm that potentiation within the SC was causing CS-evoked dopamine release, SCH23390 (which blocked the nVEP potentiation in the SC) was injected locally into the deep layers of the SC. We found that SCH23390 significantly attenuated dopamine release elicited by visual stimulation following pairing (23.7 ± 7.9% compared to control), but left intact dopamine release in response to electrical stimulation of the SNc/VTA during pairing (95.9 ± 13.1% compared to control; **Fig.** 5b,c, Extended Data Fig. 5).

**Fig. 5.**
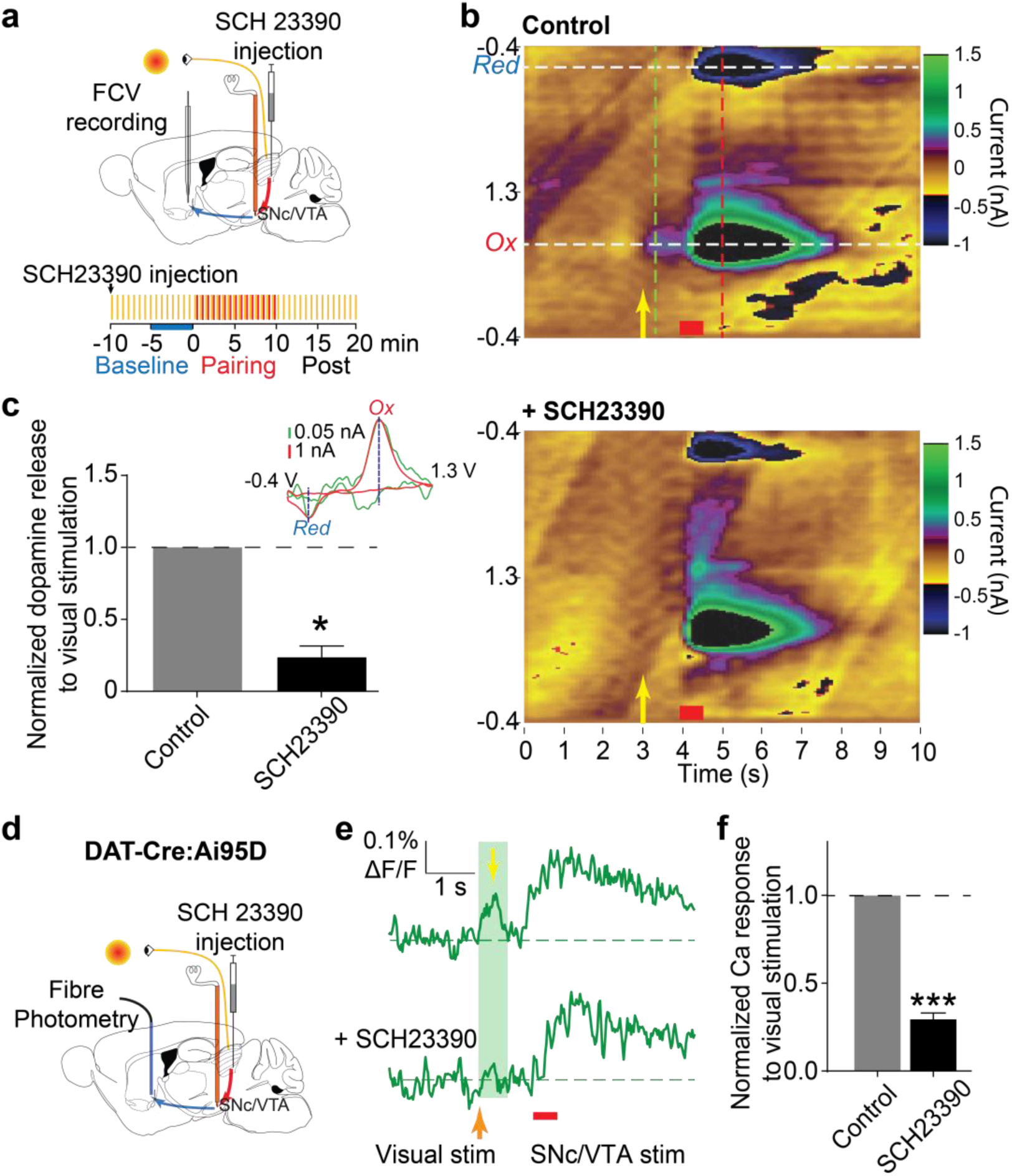
SC drives dopamine release in the striatum through classical conditioning. **a**, Striatal dopamine release via activation of the tectonigral (red arrow) to mesostriatal/mesoaccumbens (blue arrow) pathways was disrupted by local injection of D1 antagonist, SCH23390, in the deep layers of SC. **b**, A representative colour plot of FCV recording during the pairing of visual (yellow arrow) and electrical stimulation (red bar) before (upper) and after (lower) SCH23390 injection. Black indicates saturated values in small scales (see Extended Data Fig. 5 for normal scale). **c**, Example voltammograms following visual (green) and electrical (red) simulation (inset). SCH23390 blocks dopamine release following visual stimulation (via tectonigral to mesostriatal/mesoaccumbens pathway) but not electrical (mesostriatal/mesoaccumbens pathway alone) stimulation (One-sample t-test, *P < 0.05, N = 3). **d**, fibre photometry recording of GCaMP6f signal in NAcc. **e**, Examples of Ca^2+^ imaging responses in dopamine axons in NAcc. The yellow arrow indicates visual response. **f**, local injection of SCH23390 in SC blocked dopamine axon responses to visual stimulation in the NAcc (One-sample t-test, * P < 0.05, N = 4).

Additionally, we measured dopamine axon activity in the NAcc by imaging calcium signals, which are more readily detectable than dopamine by FCV and do not require cocaine to amplify the signal (**Fig. 5d**). Fibre photometry recording of GCaMP6f in NAcc in DAT-Cre:Ai95D mice showed again that local application of SCH23390 in the deep layers of the SC significantly attenuated dopamine axon activity in the NAcc following visual CS (30.6 ± 2.0% compared to control; **Fig. 5e&f**).

Notably, the dopamine release and calcium signal in dopamine axons recorded in NAcc are most likely triggered by dopamine cell body activity. Cholinergic interneurons, as a possible alternative mechanism, are too slow to drive^23^ or may even prevent^24^ dopamine axon activity.

Therefore, dopamine-dependent potentiation of the nVEP in the deep layers of the SC is able to drive dopamine release into the striatum following CS.

## Discussion

We have demonstrated that the deep layers of the SC identify the salience of the visual CS. The potentiated response developed in the deep layers of SC can drive the midbrain dopamine neurons to release dopamine in the striatum at the latency of visual classical conditioning. Many brain areas that have inputs to the midbrain dopamine neurons have been discovered to carry information about classical conditioning ^25^. However, the brain structure where the response to the CS originates is still unidentified. Our study, for the first time, confirms that the deep layers of the SC form an association between a CS and US during visual classical conditioning and drive downstream dopamine responses.

Understanding how activation of the dopamine neurons following a US modulates their own firing response to a CS alone is critical for understanding the processes of classical conditioning. Our results suggest that the dopamine signal that accompanies a US potentiates the neuronal response in SC elicited by a CS such that a CS alone can drive the dopamine neurons to release dopamine in target areas. By introducing this feed-forward loop into the circuit, our model (**Fig. 6**) provides a possible explanation of the modulatory effect of dopamine in classical conditioning, at least for visual stimuli.

**Fig. 6.**
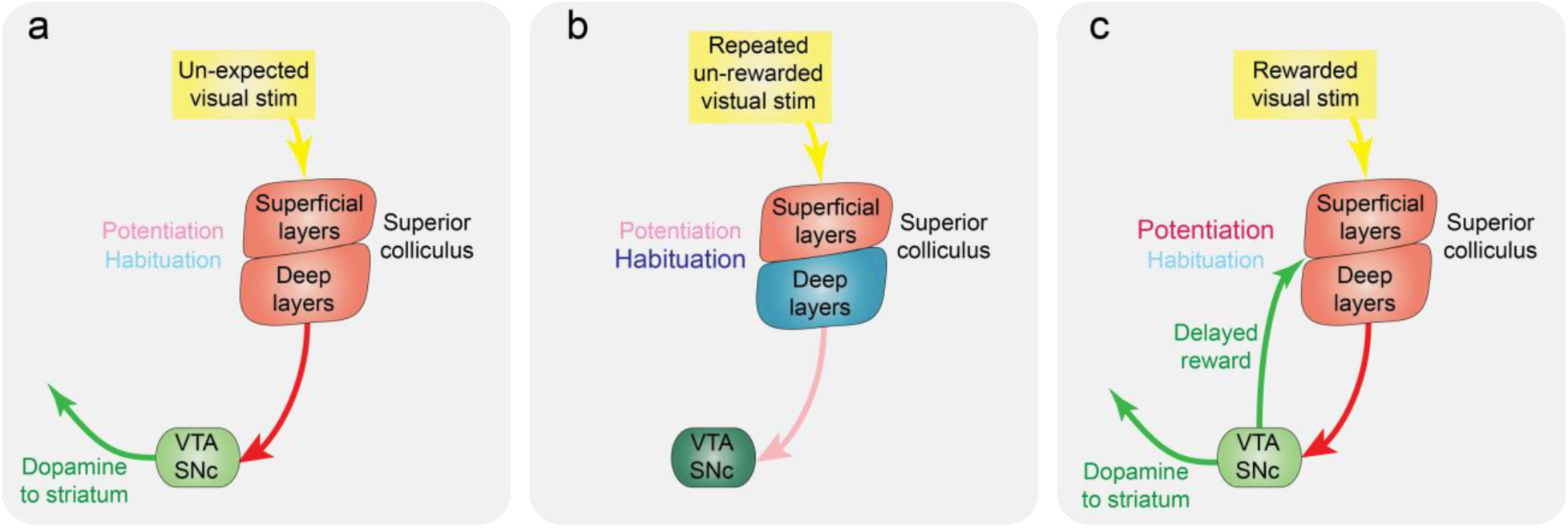
A model for how the deep layers of SC drive striatal dopamine release during stages of visual classical conditioning. **a**, A salient un-expected visual stimulus drives the deep layers of the SC to activate dopamine neurons in SNc/VTA. **b**, When the visual stimulus loses its salience, the deep layers of the SC habituate to the stimulus. **c**, The delayed dopamine signal elicited by the US potentiates the visual response and the SC starts to drive dopamine release into the striatum following visual stimulation (CS). The balance of habituation and potentiation, therefore, controls visual responsiveness in the SC.

In the current study, we applied electrical stimulation to electrodes in the dopamine cell layer (**Extended Data Fig. 5**) and used an electrically-activated dopamine signal as the US to precisely control the interval of CS and US. Since there was no requirement for any movement by the animals to receive the reward, this paradigm was suitable for study in both anaesthetised and freely moving animals. We showed that the nVEP in the SC can be potentiated when a visual stimulus is followed by a reward 1 s later, which is consistent with action discovery paradigms in humans and animals ^26, 27^. However, it does not imply that the SC can only form conditioned associations when rewards are delivered within a critical 1 s to 2 s delay. The 1 to 2 s optimal reward delay observed in our experiment might be specific for using SNc/VTA stimulation as a reward, which only paired with the visual stimuli 60 times. Also, many of our results were acquired with anaesthetised animals. It is possible that in awake behaving animals, longer reward delays may be bridged by behaviorally relevant inputs to SC from cortex.

Moreover, it is possible that longer reward delays (>2 s) could potentiate the nVEP after more pairings ^2, 28^ than the 60 pairings we used here.

We also found that a D_1_ receptor antagonist delivered to the SC blocked the development of the nVEP. In addition to the dopamine neuron projection from the midbrain to mid and deep layers of SC ^29^, the SC receives dopamine innervation from A13 zona incerta ^30^. In addition, all layers of the SC receive massive serotonergic projections from the raphe nuclei ^31, 32^, and express dense 5-HT_1A_ and 5-HT_2_ receptors ^33^. Here we show that 5-HT_1A_ but not 5HT- _2A_ receptors are involved in forming an association between CS and US in deep layers of SC. While blocking 5-HT_1A_ receptors reduces the excitatory effect of optic tract stimulation on SC neurons ^33^, and ketanserin (a 5-HT_2_ antagonist with primary action at 5-HT_2A_ receptors) attenuates the excitatory effects of 5-HT on SC neurons^34^, our results suggest that serotonin is involved in associating visual SC and US by modulating the excitatory effect of the optic input onto SC neurons.

Dopamine neurons can behave in a similar pattern when responding to a CS during classical conditioning^35^, making it possible that they are driven by a common input. Neurons in many brain areas sending input to dopamine neurons have been shown to encode the saliency of sensory cues in classical conditioning^36, 37^. While dopamine neurons increase their firing rate in response to a CS, this response is likely caused by excitatory input to dopamine neurons.

Here, our data suggest that the deep layers of the SC drive dopamine neurons to respond to a CS in classical conditioning via their excitatory glutamatergic input. This conclusion supports a recent finding that glutamatergic input may drive dopamine phasic activity in response to a CS^38^.

Our research not only provides new insights into the functioning of the SC, but also has practical implications. The potentiated neuronal response in the SC not only drives the dopamine neurons during classical conditioning, but also provides a physiologically meaningful measure of the ‘eligibility’ trace. This trace, a short-lasting neural effect induced in the SC by the light flash, is operated on by reward within a critical timing interval (less than 2 seconds). By measuring the neuronal activity, we can characterise the eligibility trace for visual stimuli within SC. This method could be extended to characterise the eligibility trace for other sensory modalities and using other behavioural paradigms, potentially leading to new approaches in the field of neuroscience and psychology.

We note that our experimental paradigm is unable to distinguish the different components of the dopamine response to the CS. It has been demonstrated that dopamine neurons can respond to a CS with two distinct components when the CS is sufficiently complex, such as when it includes information about the probability of receiving a reward. While the first large component encodes the novelty of the CS, the smaller second component can encode the reward probability^39^. In our experiment, a simple light flash was used as the CS, and it indicates a 100% probability of reward during the pairing of CS and US. Therefore, the potentiated neuronal response in the deep layers of the SC we recorded may only contribute to dopamine response component that encodes the novelty of the CS.

In summary, we propose that early sensory processing in the deep layers of the SC encodes the value of visual stimuli via the process of classical conditioning. The visual response in this structure represents a dynamic balance of potentiation, elicited by a dopamine/serotonin signal in the SC driven by the US (directly or indirectly), and the properties of habituation, which by default attenuates the response to unreinforced stimuli (**Fig. 5**). This mechanism of sensory conditioning would ensure that the system remains dynamically responsive to biologically significant sensory events. Importantly, it offers a potential explanation of not only the extensively reported sensory response characteristics of dopamine neurons ^16, 40^, but also the enhanced ability of biologically salient stimuli to attract attention ^41–43^.

## Acknowledgements

We thank Dr. Sarah Threlfell, Dr. Katie Jennings, Dr. Katherine Brimblecombe and Dr. Nicolas Vautrelle for helpful discussions and Julia Jenkins for assistance with TopScan behavioural analysis. We are very grateful to Profs. Okihide Hikosaka, Peter Dayan, Minoru Kimura and Dr Anushka Fernando for their critical comments on an earlier version of the manuscript. We thank Dr. Mark Walton and Dr. Clio Korn for their assistance with *in vivo* FCV recordings. Y.F.Z. received a Postgraduate Scholarship from the Department of Anatomy at the University of Otago. This work was funded by a grant from the Marsden Fund of the Royal Society of NZ (J.N.J.R., W.C.A. and P.R.), Academy of Medical Sciences Springboard award supported by the British Heart Foundation, Diabetes UK, the Government Department for Science, Innovation and Technology (DSIT) and Wellcome (SBF009\1125, Y.-F. Z). a grant from Parkinson’s UK (Y.-F.Z. and S.J.C.), and a grant from the Wellcome Trust (213465, A.L.).

## Contributions

Y.-F.Z. designed experiments, collected and analysed data, and drafted the manuscript. J.D. collected and analysed multi-unit recording data. M.J.B. performed some of the histology and provided technical support. S.J.C. provided technical support for in vivo FCV. E.M., W.C.A., and P.R. contributed to experimental design and data interpretation. All authors critically reviewed the manuscript. J.N.J.R. designed experiments and reviewed the manuscript. Y.-F.Z. and J.N.J.R. supervised the study.

