## Extended Data Figs for "The superior colliculus gates dopamine responses to conditioned stimuli in visual classical conditioning"

**This PDF file includes:**

Materials and Methods
Extended Data Fig1 to 6

### Materials and Methods

All experimental procedures with rats were approved by the University of Otago's Animal Ethics Committee in accordance with the NZ Animal Welfare Act 1999. All procedures with mice in FCV and multi-unit experiments were performed in accordance with the UK Animals (Scientific Procedures) Act 1986 (Amended 2012) with ethical approval from the University of Oxford and under the authority of a Project Licence granted by the UK Home Office.

#### Surgery for recording with naïve rats

A total of 68 male Long Evans rats weighing 350 to 390 g were anesthetised with an initial dose of urethane (1.6-2.0 g/kg, Sigma-Aldrich, intraperitoneal; i.p.) and an additional dose 15 minutes later (0.6 g/kg, i.p.). Before surgery, a local anaesthetic (bupivacaine, 0.5%) was injected into the scalp. Supplementary doses, 0.2 ml, of urethane (0.6 g/kg) were given through an i.p. cannula during experiments to maintain the anaesthesia level. Core temperature was maintained at 35-36°C during the experiment using a homoeothermic blanket and rectal probe (TR-100, Fine Science Tools). A concentric stimulating electrode (Rhodes NEW-100X 10 mm, USA) was implanted through a skull hole into the SNc/VTA in the left hemisphere (AP -4.8 mm and ML 1.5 mm to Bregma, at a depth of 7.7-7.8 mm from the brain surface). Electrode positions were verified in cresyl violet stained sagittal sections (**Extended Data Fig 6**).

#### Surgery for recording multi-unit activity with mice

C57Bl/6 mice were anaesthetised with an i.p. injection of urethane (1.5 g/kg). A local anaesthetic (0.5% Bupivacaine) was injected into the scalp before surgery. Core temperature was maintained at 36°C during the experiment using a homoeothermic blanket and rectal probe (Harvard Apparatus). Eye lubricant was applied at the beginning of the surgery, and the light source was placed ca. 1cm from the contralateral eye. The stimulation electrode was placed in the SNc/VTA (AP -3.1, ML 0.8, DV -4.2), and the recording electrode was placed in the lower SC (AP -4.0, ML 0.8, DV -2.1). Silicon 15-channel electrodes (Cambridge Neurotech) were used, with a triangular electrode arrangement at the tip spanning 150µm vertically. A reference wire was introduced into the contralateral cerebellum.

**Behavioral methods**

Seven male Long Evans rats (350±390 g) were implanted for ICSS. Rats were anaesthetised with ketamine (60 mg/kg i.p., Phoenix Pharm Distributors Ltd, NZ) and domitor (0.5 mg/kg, i.p., Orion Pharma Pfizer, Finland) and placed in a stereotaxic frame. Prophylactic antibiotic was administered subcutaneously. A bipolar stimulating electrode (PlasticsOne MS303/2, USA) was positioned at the level of the SNc/VTA as in naïve rats. At least three days were allowed for post-operative recovery. Rats were trained to perform ICSS in a behavioural chamber with a lever that, when pressed, immediately triggered delivery of a single stimulus train (100 Hz, 0.5 ms biphasic pulses, 500 ms train duration) to the SNc/VTA electrode. The response rate versus current intensity data was collected daily. The training was ceased when the optimal current was stable to within 10% for three consecutive sessions (Fig. 4a).

In behavioural conditioning experiments, the animals were placed in the recording chamber and allowed to settle for 15 minutes in the dark. A program written in MedPC (Med Associates, USA) then began to deliver a short activation (30 ms) of five stimulus lights around the chamber, ganged to achieve whole field visual stimulation, at an interval of 10 seconds. After a 10-minute baseline period of light flashes alone, each light presentation was followed 1 s later by the delivery of a single stimulus train to the animal's SNc/VTA at their optimal current. The LFP in the deep layers of the SC was recorded for offline analysis. The behavioral response to the light was video recorded throughout the experiment and analysed offline using TopScan behavioral analysis software. The average speed of the animal's movements in 0.5 s bins around each light flash was measured and exported to Excel files. The difference in speed 0.5 s before and after each light flash was calculated during the pairing protocol and during the baseline period where the light flash was delivered alone prior to pairing. The difference in speed between 0.5 s after the SNc/VTA stimulation and the period prior to the light flash during pairing was also calculated for comparison.

**Local field potential recording**

The LFP in the superior colliculus was recorded using a glass electrode (1-2 MΩ) pulled from calibrated glass capillaries (volume 5 µl, diameter 1.0 mm; Modulohm I/S,

Denmark) filled with 0.9% NaCl solution. The recording electrode was implanted in the SC (Fig. S6b,c) with a micromanipulator (IVM, Scientifica, UK) at the position of AP -6.5 mm and ML 1.5 mm, or -1.5 mm when recorded from both hemispheres, to Bregma, at a depth of 4.0-4.1 mm from the brain surface. The coordinates were chosen to maximise the possibility of recording the LFP from the midpoint of the deep layers of the SC in the mediolateral plane, so the LFP did not only reflect the properties of medial or lateral SC[44]. A Teflon-coated tungsten wire was immersed in the NaCl solution, and connected via a headstage (NL100 Neurolog) to a preamp (NL104), an amplifier (NL106) and a filter (NL125). The animal was grounded by a silver wire, which was introduced into the connective tissue underneath the animal's back skin. Both signals were amplified and band-pass filtered (0.1 to 10,000 Hz). All waveform data were digitised at 50 kHz by 1401 Micro 2 (CED), displayed with Spike2 software (CED) and stored to disk. Behaving animal recordings were performed using a chronically implanted stainless steel wire and reference electrode screwed on the skull on the other side above the SC connected to a locally-constructed battery-powered FET headstage, run with the stimulator wires to a commutator (PlasticsOne, USA) and then to a preamplifier and Neurolog recording system.

**Multiunit activity recording**

Recordings with the Silicon 15-channel electrodes (Cambridge Neurotech) electrode were acquired using Intan Technologies data acquisition software (RHX) at 30kHz, and the unfiltered data was stored to disk. Data analysis was done using custom-written MATLAB and Python scripts. Spikes were extracted from the unfiltered data using the Kilosort v2.0 software. All spikes on each channel were then pooled to represent the multi-unit activity. The multi-unit activity (MUA) response magnitude was calculated as the difference between the spike rate 50-100ms after light stimulation and the baseline spike rate (from 1 second before light stimulation).

**Drugs**

Drugs were injected through the glass recording electrode in the SC. The pipette was held by a pipette holder that was attached to polyethylene tubing with a 10 ml syringe as a pressure injector. Each drug, including BIC (0.01% in saline, 250 nl), SCH23390 (2 µg in 250 nl saline), WAY100635 (2 µg in 250 nl saline), ketanserin (2.5 µg in 500 nl saline), or saline (250 nl), was injected into the SC at a rate of 400 nl/min. All drugs were sourced from Sigma-Aldrich.

**Light and electrical stimulation**

Light stimuli (10 ms duration, 0.1 to 0.033 Hz) were delivered by a white LED (1500 mcd) that was placed 1-2 cm directly in front of the right eye of the animal. The left eye was covered. In eight experiments, stimuli were delivered to both eyes alternately with the non-stimulated right eye covered. In the experiments with naive rats, SNc/VTA stimulation (0.5 ms dmiphasic pulses, 500 ms train duration; no damage noted in any experimental brain sections) was delivered at 100 Hz at a current of 500 µA. In the experiments with ICSS rats, SNc/VTA stimulation (100 Hz, 0.5 ms biphasic pulses, 500 ms train duration) was delivered using the optimal current acquired during training. Stimulation was controlled by locally developed software “StimulatorControl” (SCL Limited, New Zealand) on a Windows-compatible PC computer.

**Analysis of electrophysiological recording data**

Data were analysed using Spike2 v6.09 and MATLAB 2012a, including customised MATLAB scripts. The average LFP was generated with a built-in function of Spike2. The onset time of visual stimulation was used as a trigger to create the average of the LFP over a 5-minute recording (30 trials). The 50 ms period prior to the visual stimulation was defined as the baseline of the trace, and its average value was defined as 0 mV. The amplitude of the negative component of VEP (nVEP) was defined as the minimum value of the LFP during the period of 60ms to 100ms, followed by the visual stimulation (Fig. 1b). To minimise the influence of the recording noise, the amplitude was read as the average of eleven data points around the minimum value of the LFP.

Because the nVEP showed its minimum value at the last 5 min of baseline recording and its maximum value during the last 5 min of pairing, the difference of the average LFP between these two periods was defined as the amplitude of the nVEP (Fig. 1).

***In vivo* fast-scan cyclic voltammetry (FCV)**

A total of 16 adult (10 - 12 weeks) male C57Bl6/J mice were used for the in vivo FCV experiments. Mice were anaesthetised with urethane (1.5 – 1.8 g/kg), and surgical anaesthesia was maintained with supplementary isoflurane (1% w/o). Body temperature was maintained at 36-37 °C using a homoeothermic heating blanket. Corneal dehydration was prevented with eye ointment (Lacri-lube). After induction, the mouse was secured in a stereotaxic frame, the scalp was shaved and cleaned with dilute hibiscrub and 70% alcohol. A local anaesthetic, bupivacaine, was injected s.c. into the incision site. After the skull was exposed, holes were drilled for the Ag/AgCl reference electrode (AP: -2.0 mm, ML: 0.5 mm).

Extracellular DA concentration was monitored in the ventral striatum using FCV with 7 µm diameter carbon fibre microelectrodes (CFMs; tip length 50-100 µm). The scanning voltage was a triangular waveform (-0.4 to +1.3 V range vs Ag/AgCl) at a scan rate of 400 V/s and sampling frequency of 10 Hz, which was produced from a Tarheel system (USA). The data was acquired and analysed offline with custom-written Matlab (R2013b) scripts.

#### *In vivo* Fiber Photometry

Round pieces of skull overlying the left hemisphere were removed to allow access to the DLS (AP +1.0 mm, ML 1.6 mm, DV 2.2 mm to Bregma) and SNc (AP -3.1mm, ML 0.8mm, DV 4.3mm to Bregma). The injection and recording array, consisting of a glass pipette and a 200 µm diameter fibre, was positioned in the DLS. GCaMP6f expressed in DA axons in DAT-Cre:Ai95D mice was activated with 480 nm light (76 µW), and the intensity of GCaMP6f emission was sampled at 40 Hz with Neurophotometrics (FP3001).

For midbrain electrical stimulation, 0.5 mA current (500 µs) was delivered through a bipolar stimulating electrode (0.005 inch, MS303/3-A/SPC, P1 Technologies) at 0.1 Hz. The stimulating electrode tips were separated by ~500 µm.

#### SNc/VTA Stimulation and drug injection during in vivo FCV

A bipolar stainless steel stimulating electrode and a drug injection pipette pulled from a calibrated glass capillary (volume 5 µl, diameter 1.0 mm; Modulohm I/S, Denmark) were implanted in the SNc/VTA (AP -3.5 mm, ML -0.35 mm, DV 4.3 to 4.7 from brain surface) and SC (AP -3.5 mm, ML -1.0mm, DV 1.8-2.2 mm from brain surface), respectively. SNc/VTA stimulation (0.5 ms biphasic pulses, 1200 ms train duration) was delivered at 50 Hz with the current at 500 µA. Note the stimulation frequency was chosen to avoid the clash of stimulation artefacts with FCV recording sweeps.

### Experiment protocol

The LFP recording in the SC was similar for all anesthetized animal experiments:

1) Baseline recording: After the surgery, the rat was covered and kept within the dark for at least 10 min, so the animal adapted to the dark environment. Once the LFP stabilized, i.e. the LFP showed slow synchronized oscillation, the visual stimulation was applied to the right eye of the animal at 0.1 Hz for 10 min.
2) Pairing protocol: The pairing of visual stimulation and SNc/VTA stimulation was applied at 0.1 Hz for 10 min.
3) Post-pairing protocol: The visual stimulation was applied at 0.1 Hz for 10 min.

Drug injection: Drugs were injected directly before the baseline recording.

For each animal, at most two experiments were applied. There was a 30 min break between the experiments. During the break, no stimulation was applied to the animal.

### Histology methods

When the recording experiment was finished the animal was perfused with a 0.1 M phosphate buffer solution (PBS) containing 4% paraformaldehyde. The brain was then removed and kept in fixative solution at 4°C. Brains were sliced in 60 µm sections, which were then stained with a 0.03% cresyl-violet solution, and the correct position of the tracks was confirmed under the light microscope.

### Statistics

Analyses of the nVEP were carried out using one-way ANOVAs with Bonferroni's Multiple Comparison post hoc test, Friedman tests (repeated, nonparametric) with Dunns post hoc test, or Paired t-tests. Analyses of dopamine release in the striatum were carried out using a one-sample t-test. Statistical analyses were performed using Prism5. Data are mean ± S.E.M. unless stated. Significance levels are indicated as follows: * $P < 0.05$; ** $P < 0.01$; *** $P < 0.001$; **** $P < 0.0001$.

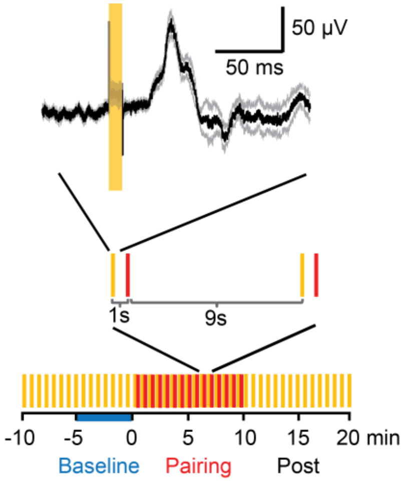


**Extended Data Fig. 1.** A diagram of the typical experimental protocol. Visual stimulation (orange) was applied every 10 s. To potentiate the short latency (60 – 100 ms) visual response in the deep layer of the SC (black and grey line, upper panel), electrical SNc/VTA stimulation (red) was applied 1 s after the visual stimulation, 60 times. Variations of these timing relationships were tested in some experiments.

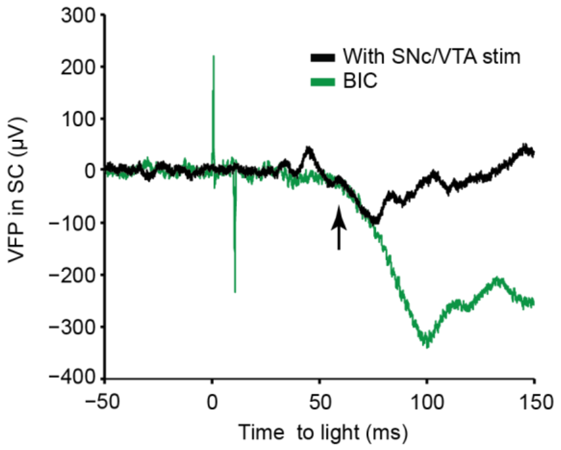


**Extended Data Fig. 2** The VEP induced by visual classical conditioning shares similarities with the VEP induced by local injection of BIC in one rat. The potentiated nVEP component induced by pairing with SNc/VTA stimulation (black; pairing minus baseline trace) occurs at a similar onset latency (arrow) but earlier peak latency to the nVEP under the disinhibitory effect of BIC (green).

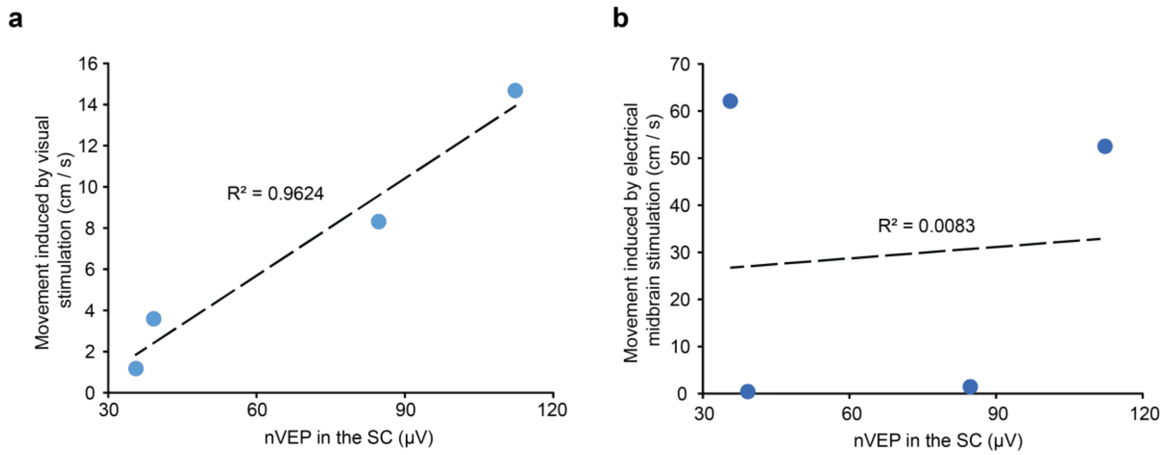


**Extended Data Fig. 3** The development of nVEPs in the deep layers of SC correlated with the learning of the salience of the CS. **a**, The difference in the speed of movements in response to the light between the pairing protocol and the pre-pairing baseline period was positively correlated with the pairing minus baseline amplitude of the nVEP in the deep layers of SC. **b**, The difference in movement speed in response to the electrical stimulation was not correlated to the nVEP. Thus the change in the nVEP through pairing was related to the learning of the salience of the visual stimulus (CS; as indicated in a) and not to a general behavioral activation induced by the electrical stimulation (US).

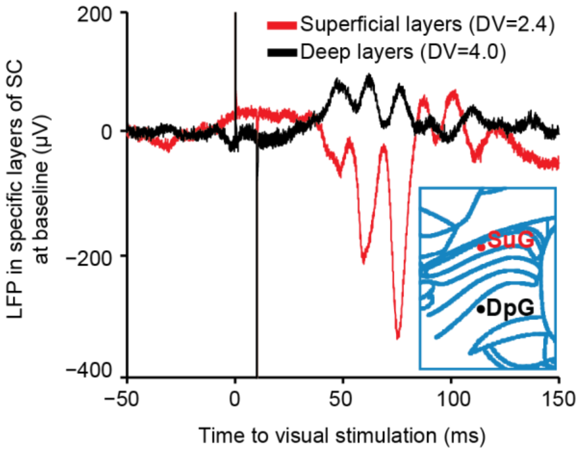


**Extended Data Fig. 4** The mean VEP in the deep layers (black) and the superficial layer (red) of the SC, recorded in the same animal using the same electrode, before pairing. The positive component of the VEP in the deep layers is aligned in time with the negative component of the VEP in the superficial layers, indicating that pVEP in the deep layers represents the current sink for the nVEP in superficial layers of the SC. SuG: superficial grey layer of the SC, DpG: deep grey layer of the SC.

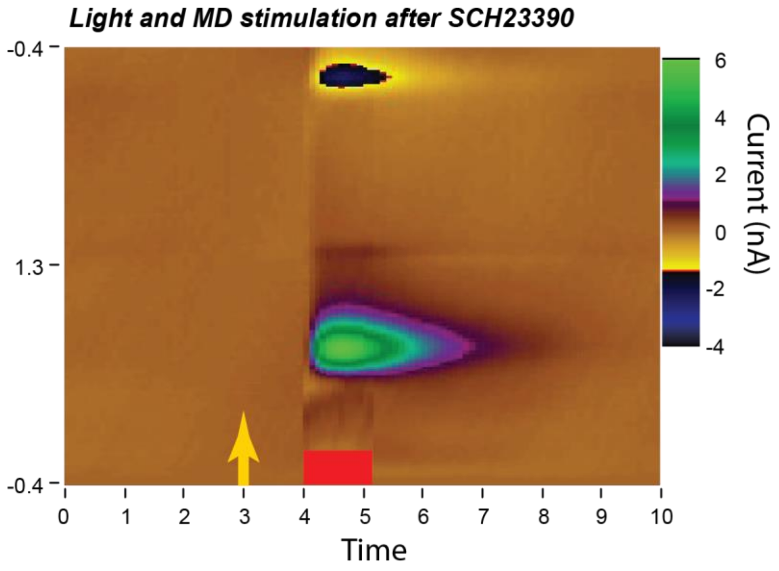


**Extended Data Fig. 5** Dopamine release in response to the electrical stimulation during the pairing protocol on a normal scale. Local injection of SCH23390 blocks the dopamine release following visual stimulation, as in Fig. 5b. The scale is set to usual values to correctly show the dopamine release following midbrain dopamine (MD) electrical stimulation (red bar).

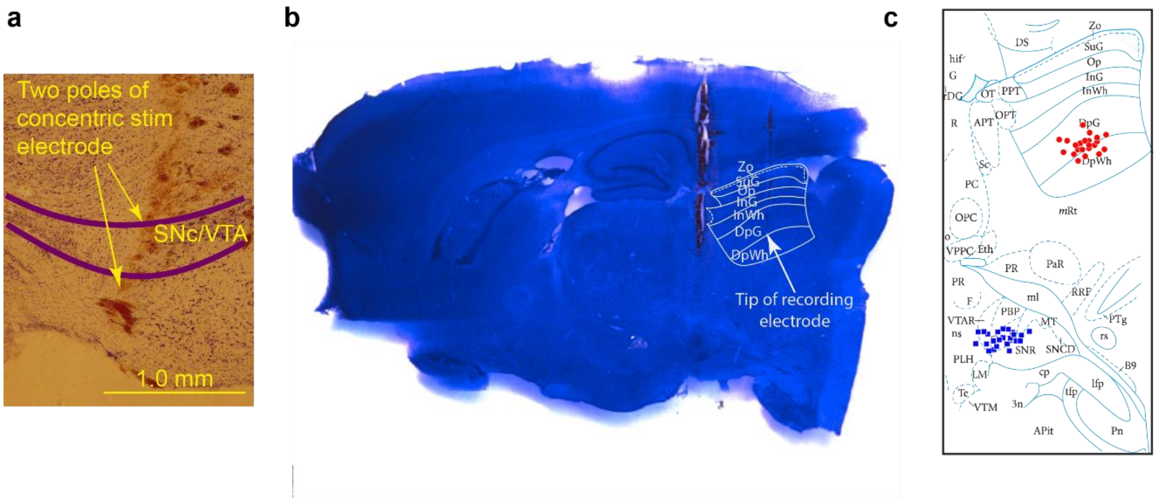


**Extended Data Fig. 6** Example histological sections showing stimulating and recording electrode locations. **a**, The concentric stimulating electrode positioned to activate SNc/VTA. **b**, the tip of the recording electrode is positioned in the middle of the deep layers of the SC. Part of the stimulating electrode can also be seen more anteriorly in this sagittal section, in its path to the midbrain. DpG: deep grey layer, DpWh deep white layer of the SC. **c**, Approximate midpoint of the stimulating electrodes (blue squares, sagittal section at mediolateral +1.5 mm) and the position of recording electrode tips (red circles).
